# Principles and characteristics of biological assemblies in experimentally determined protein structures

**DOI:** 10.1101/564385

**Authors:** Qifang Xu, Roland L. Dunbrack

**Affiliations:** Institute for Cancer Research Fox Chase Cancer Center 333 Cottman Ave. Philadelphia, PA 19148, USA

## Abstract

More than half of all structures in the PDB are assemblies of two or more proteins, including both homooligomers and heterooligomers. Structural information on these assemblies comes from X-ray crystallography, NMR, and cryo-EM spectroscopy. The correct assembly in an X-ray structure is often ambiguous, and computational methods have been developed to identify the most likely biologically relevant assembly based on physical properties of assemblies and sequence conservation in interfaces. Taking advantage of the large number of structures now available, some of the most recent methods have relied on similarity of interfaces and assemblies across structures of homologous proteins.

## Introduction

Proteins function by their interactions with other molecules, including other proteins, nucleic acids, and small molecules. Many and perhaps most proteins function as homo-and heterooligomers, containing two or more copies of at least one subunit type. The structures of these protein complexes are referred to as *biological assemblies*. Structure information on biological assemblies is obtained by X-ray crystallography, nuclear magnetic resonance, cryo-electron microscopy, and other experimental methods. In this review, we discuss the principles and characteristics of biological assemblies. We first define several of these principles and characteristics and then examine recent developments relevant to each of these areas. We discuss recent computational approaches for identifying the correct assembly present in a crystal and how these methods should be benchmarked.

### Principles of biological assemblies in experimental protein structures

1. A **biological assembly** is a functionally relevant complex of proteins and perhaps other molecules. It can be defined structurally by its Cartesian coordinates and by its stoichiometry, molecular interfaces, and symmetry. The biological assembly of an experimental structure is defined as the largest functional form present in the experimental data. This assembly is usually one of the functional forms found in vivo, although in some cases it may not be since experimental conditions or protein constructs may affect the oligomeric state of a protein in experimental structure determination. For cryo-EM, the structure of the biological assembly is obtained more-or-less directly from the experiment [1]. For NMR, the experimental data may provide distance constraints between residues in different subunits and thus provide direct information on contacts needed to build an assembly. For crystal structures, the biological assembly is either the asymmetric unit, a subset of the asymmetric unit, or parts or the whole of multiple asymmetric units from a single unit cell or multiple neighboring unit cells. The biological assembly of a crystal structure is often ambiguous. The PDB contains author annotations of biological assemblies in the form of Cartesian coordinates, which for crystal structures differ from the asymmetric unit (ASU) about 40% of the time. Typically, crystals contain one assembly type, as defined by stoichiometry, interfaces, and symmetry, that covers the entire crystal (e.g., a crystal cannot be a lattice of dimers of one type and monomers that do not form the dimer).
2. **Stoichiometries** of biological assemblies define the number of each subunit type (designated by Roman letters) in the assembly (usually denoted A, A2, A3, AB, A2B2, etc.) and may be even (same number of each subunit type: AB, A2B2C2, etc.) or uneven (different numbers of different subunit types: A2B, A3B2, etc.). For crystal structures, ordinarily the biological assembly should have at least one copy of every protein entity (sequence) in the asymmetric unit, although some entries are incorrectly annotated and missing some subunit types in the first biological assembly.
3. **Interfaces** between pairs of proteins in biological assemblies can be isologous (homodimeric and symmetric) or heterologous (homodimeric or heterodimeric, and asymmetric) or pseudoisologous (heterodimeric but approximately symmetric interfaces of homologous protein domains, as determined by sequence and/or structural alignment). Similar interfaces are typically present across multiple crystal forms of the same or related proteins when all or part of the assemblies are also conserved. Biologically relevant interfaces are typically larger and more conserved in sequence than interfaces induced by crystallization. However, some assemblies held together by large, primary interfaces may also contain some small, non-conserved interfaces that are induced by assembly formation of the primary interfaces (e.g., a cyclic tetramer may have one large interface that occurs four times but a small one along the diagonal).
4. **Symmetry** types of biological assemblies observed in the PDB include asymmetric (C1), cyclic (Cn, n≥2), dihedral (Dn, n≥2), icosahedral, helical, tetrahedral, and octahedral. Cn assemblies have a single *n*-fold axis of symmetry. C2 cycles contain an isologous interface. Proteins in C3 and larger cycles are connected by heterologous interfaces around the cycle. Dn assemblies contain two symmetry axes: they consist of C2 symmetric assemblies of two Cn cycles. The interfaces between the cycles are all isologous. Symmetry can be global -- covering the whole assembly, or local -- covering only a subset of subunits in the assembly. Symmetry can be exact – involving copies of the same subunits related by translations and/or crystallographic symmetry operators, or approximate – involving different monomers from the asymmetric unit of the crystal. Most assemblies are “closed,” that is not extending indefinitely within a crystal lattice, with the exception of helical or filamentous assemblies found in some structures, such as actin.
5. **Benchmarking of computational methods** for determining the biological relevance of interfaces or assemblies should be based on a comparison of the Cartesian coordinates of the predicted interfaces and assemblies and the “true” interfaces or assemblies. However, the most commonly used benchmarks list the stoichiometry and symmetry of the supposedly correct assemblies without defining the coordinates. There are numerous discrepancies between the symmetries and stoichiometries of the “true” assemblies in these benchmarks and the biological assemblies deposited for these entries in the PDB. Even when the symmetry and stoichiometry of a predicted assembly agree with those of the deposited assembly, it is not guaranteed that the deposited assembly is correct (e.g., a crystal can contain more than one C2 homodimer that may be a plausible biological assembly).
6. **Methods for distinguishing biologically relevant interfaces** (i.e., present in biological assemblies) from crystal-induced interfaces are based on biophysical properties of the interfaces, sequence conservation of amino acids in the interfaces, and/or structural conservation of the interface across multiple crystal forms of the same or related proteins. An analysis of structural conservation across multiple PDB entries provides more information when crystallographic space groups are used to build all possible interfaces within the crystal, rather than depending only on those in the asymmetric unit or the deposited or predicted biological assemblies.
7. **Methods for identifying the correct biological assembly** within crystals are based on many of the same criteria that authors use (physical properties, symmetry, sequence, and structural conservation). They also depend on dividing a crystal into possible assemblies based on the principle of closed assemblies and that a crystal should have only one form of assembly that covers the whole crystal. They can be more successful if they consider all possible assemblies within a crystal, not just those that have been deposited as such in the PDB or predicted by other programs.
8. **There are exceptions to every rule…** We discuss each of these areas in turn.

### Biological assemblies

Crystallographic data analysis provides the asymmetric unit (ASU), which can be used to build a model of the crystal lattice via symmetry operations. However, from the earliest days of protein crystallography, it was realized that biologically relevant assemblies may not necessarily coincide with the ASU. After the first structure of myoglobin was determined by Kendrew and colleagues [2], Perutz et al. determined the structure of hemoglobin, which they observed to be a pseudosymmetric tetramer of myoglobin-like alpha and beta subunits, that could only be constructed from two copies of the alpha-beta heterodimer in the ASU (a related structure was later deposited in the PDB, now entry 2DHB) [3]. One of the earliest homooligomeric structures was that of concanavalin A with a monomeric ASU and a tetrameric biological assembly containing three different interfaces and dihedral symmetry [4].

The biological assembly present in a structure determination can be defined explicitly by the Cartesian coordinates of the assembly. These coordinates are required by the PDB during deposition of experimental structures, and are available as separate coordinate files from those of the default PDB file (the asymmetric unit for crystal structures). While the stoichiometry of the biological assembly may be known by experiments such as analytical centrifugation or dynamic light scattering, there may still be multiple choices for the correct assembly within a crystal. In many cases, the correct assembly is relatively clear – either because it is similar to previously determined structures of the same protein or homologous proteins in other crystal forms [5], or because it is far more consistent with the typical characteristics of protein oligomers (symmetry, buried surface areas, hydrophobicity, interface sequence conservation) than any other possible assembly of the correct size from the crystal [6]. Authors frequently identify a hypothetical assembly from their crystallographic data, and then test their chosen assembly in solution experiments by mutating amino acids in the assembly interface(s), although cross-linking experiments provide more informative data [7]. Often published papers provide little or no information on how the authors chose the biological assembly they deposited in the PDB. Frustratingly and quite frequently, the deposited biological assembly is not the same as shown in the paper. When this happens, the deposited assembly is usually the same as the ASU, and it seems that the authors did not appreciate the importance of depositing the correct assembly and simply copied the ASU coordinates.

The three cardinal properties of assemblies that can be derived from the Cartesian coordinates from a protein structure determination experiment are: 1) stoichiometry; 2) the interfaces and interactions that make up the assembly; and 3) the resulting symmetry of the assembly. At the protein level, for each assembly the PDB provides stoichiometries as a string of letters designating protein species and numbers indicating the number of subunits of each species. For example, A2B2 is a heterotetramer consisting of two copies of subunit type A and two copies of subunit type B. An assembly of N subunits must have at least N-1 interfaces and at most N(N-1)/2. Homodimeric interfaces within assemblies can be “isologous,” indicating that they are C2 symmetric, or “heterologous” meaning that they are asymmetric. The majority of oligomeric protein assemblies with multiple copies of one or more subunit types are symmetric. The most common symmetries are cyclic (C2, C3, C4, etc) and dihedral (D2, D3, D4, etc.), although larger symmetries occur in virus particles. The PDB provides the symmetry information of each biological assembly and the ASU.

To emphasize the difference between the ASU and the biological assembly, we compared the stoichiometry and symmetry of 144,088 entries in the PDB. The PDB does not provide biological assembly files for all NMR structures, but we can generally assume the biological assembly would be the same as the NMR ensemble PDB file. The biological assembly annotation is different from the asymmetric unit in symmetry and/or stoichiometry in 42% of crystal structures in the PDB and 38% of all entries. For 30% of PDB entries, the PDB provides more than one biological assembly. For 83% of these, the different assemblies are essentially the same, and represent different subsets of a larger asymmetric unit. But in other cases (17%), the authors or the PDB have provided hypothetical assemblies of different sizes. In some cases, the PDB provides a second biological assembly defined by the program PISA [8], especially when this assembly is larger than that provided by the authors.

Various estimates of the accuracy of the PDB’s first biological assembly exist; but values around 80% are common [5,8,9]. Fixing the obvious errors might bring this up to about 90%, while the remaining 10% may remain ambiguous because of uncertainties in the stoichiometry, interfaces, or symmetry of the true biological assembly. Some homodimers have very small interfaces (e.g. cytosolic sulfotransferases [10]), and some homodimers are formed by weak interactions that may not be present in every crystal of a protein (e.g., RAS [11]). In cases where the assembly size is not known experimentally, then the identification of the correct assembly is even more challenging.

### Stoichiometries of biological assemblies

As described above, stoichiometries are usually represented as strings of letters and counts, e.g. A4B4 is a heterooctamer of 4 copies of subunit type A and 4 copies of subunit type B. If there are more than 26 protein entities (different sequences), then lower case letter are used. If there are more than 52 entities, then the upper-case letters are used again followed by more lower-case letters. Some ribosome structures have over 80 different protein types, and hence use A-Z, a-z, A-Z, and some of a-z again. A total of 14.2% of PDB entries are heteromers (AB or larger), having two different sequences present in the first biological assembly. Stoichiometries of heteromers can be (and usually are) even, meaning an equal number of each type of subunit, e.g. AB, A2B2, A3B3C3, or uneven, meaning a different number of some subunit types, e.g. A3B2, A2BC. Of 144,088 PDB entries we evaluated, only 2,050 (1.4% of all 144,088 entries, 10% of 20,521 heteromeric entries) had an uneven stoichiometry in the first biological assembly, 513 of which were the simplest uneven stoichiometry, A2B. It is likely that some of these are not truly uneven stoichiometry, resulting from annotation errors in the biological assembly. The most common seven stoichiometries represent over 90% of PDB entries: A (monomer, 48.7%), A2 (23.6%), A4 (6.3%), AB (6.3%), A3 (2.8%), A2B2 (1.5%).

Marsh et al studied complexes with uneven stoichiometry, and found that about the same figure of 10% of heteromeric experimental structures have uneven stoichiometry, with a relative 2:1 ratio of the higher-abundance subunit(s) to the less-abundant subunit(s), representing more than 75% of structures with uneven stoichiometry [12**]. Further, protein abundance data indicate that the high-abundance subunits also have higher protein expression levels in vivo. In an analysis of 88 A2B complexes (the simplest uneven complex type), they provided a number of structural mechanisms that explain how complexes with uneven stoichiometry might form. The first is pseudosymmetry (18% of the A2B complexes) in which the B subunits contain more than one homologous domain, or repeat, each of which can bind an A subunit with similar A/B domain-domain interfaces. The second is “multi-binding” (12.5%) such that the B subunit binds the same surface region of each A subunit but on entirely different and non-homologous surfaces of the B subunit. Third (19%), a C2-symmetric, homodimeric protein (A2) can bind a third protein (B), utilizing the same surface on each A subunit to bind adjacent surface patches on B. The remaining cases involve asymmetric binding of two copies of A to B in ways that are incompatible with either A subunit binding another copy of B (i.e., to form an A2B2 tetramer), either due to steric occlusion or conformational change between the two A subunits. One caution pointed out by Marsh et al. is that 13 of the 88 structures were likely to be errors in the PDB biological assembly, as judged by the papers published on the structures [12**].

We investigated whether the rapid rise in cryo-electron microscopy is contributing significantly to the number of heteromeric structures in the PDB. Since 2010, 69% of cryo-EM structures have been heteromeric, compared to 54% in the previous decade. Conversely, 13-15% of X-ray structures and only 5-7% of solution NMR structures have been heteromeric for every decade since the 1980s. Most heteromeric structures are still solved by X-ray crystallography, but cryo-EM is catching up: in 2018, 1,423 heteromeric structures were solved by crystallography, and 597 heteromeric structures were solved by cryo-EM. However, the number of cryo-EM structures has been increasing at a rate of 30-50% per year (2,533 in 2018) while the number of X-ray crystal structures has been essentially flat for the last three years at about 9,900 per year. Cryo-EM is also providing information on oligomeric structures with repeated subunits (both homooligomers and heterooligomers with at least 2 copies of at least one protein): 68% of the 2,753 cryo-EM structures in the PDB have repeated subunits, potentially providing a rich source of interfaces for homooligomers for benchmarking biological assembly identification for crystal structures.

There are 566 structures in the PDB where the first biological assembly has a different number of protein entity types (A, B, C, etc.) than the asymmetric unit. It appears that most of these are due to inconsistent handling of peptides in the PDB in terms of stoichiometry. In some structures, the peptide is considered part of the stoichiometry, so a peptide-bound protein would have stoichiometry “AB”. In other cases, the peptide is not counted, and a peptide-bound protein would have stoichiometry “A”. There are 7 cases where the number of entity types is lower in the ASU than in the first biological assembly, because the peptide is not counted in the ASU but it is counted in the biological assembly. But there are also cases where a heterodimeric structure in the ASU is inexplicably divided into two different biological assemblies, each with one protein type. With few exceptions, these are obvious annotation errors that could be fixed in a remediation effort.

### Interfaces in biological assemblies

Monod, Wyman, and Changeux defined the properties of allosteric assemblies, including the types of interfaces found in oligomeric assemblies [13]. They defined two modes of association in terms of residue contacts: “*heterologous associations*: the domain of bonding is made up of *two* different binding sets; *isologous associations*: the domain of bonding involves two *identical* sets.” Isologous interfaces are exactly C2 symmetric if they come from two copies of the same monomer in the asymmetric unit, so that one monomer is a symmetry copy of the other. An isologous interface may be only approximately C2 symmetric if it arises from two different monomers with the same sequence from the asymmetric unit. Most contacts in an approximately isologous interface are duplicated: if [x,y] is one contact between chain 1 and chain 2 then [y,x] between chains 1 and 2 also exists. Homologous proteins with similar folds can form pseudosymmetric heterodimers, which might be referred to as pseudoisologous. Heterologous interfaces are asymmetric; it is not possible to generate one monomer from the other by a 180° rotation. Generally heterologous interfaces involve different residues on the surface of each monomer, although there can be overlap. An example of two isologous and two heterologous interfaces from a single crystal of human BRAF is shown in Figure 1 for PDB entry 1UWH [14].

**Figure 1.**
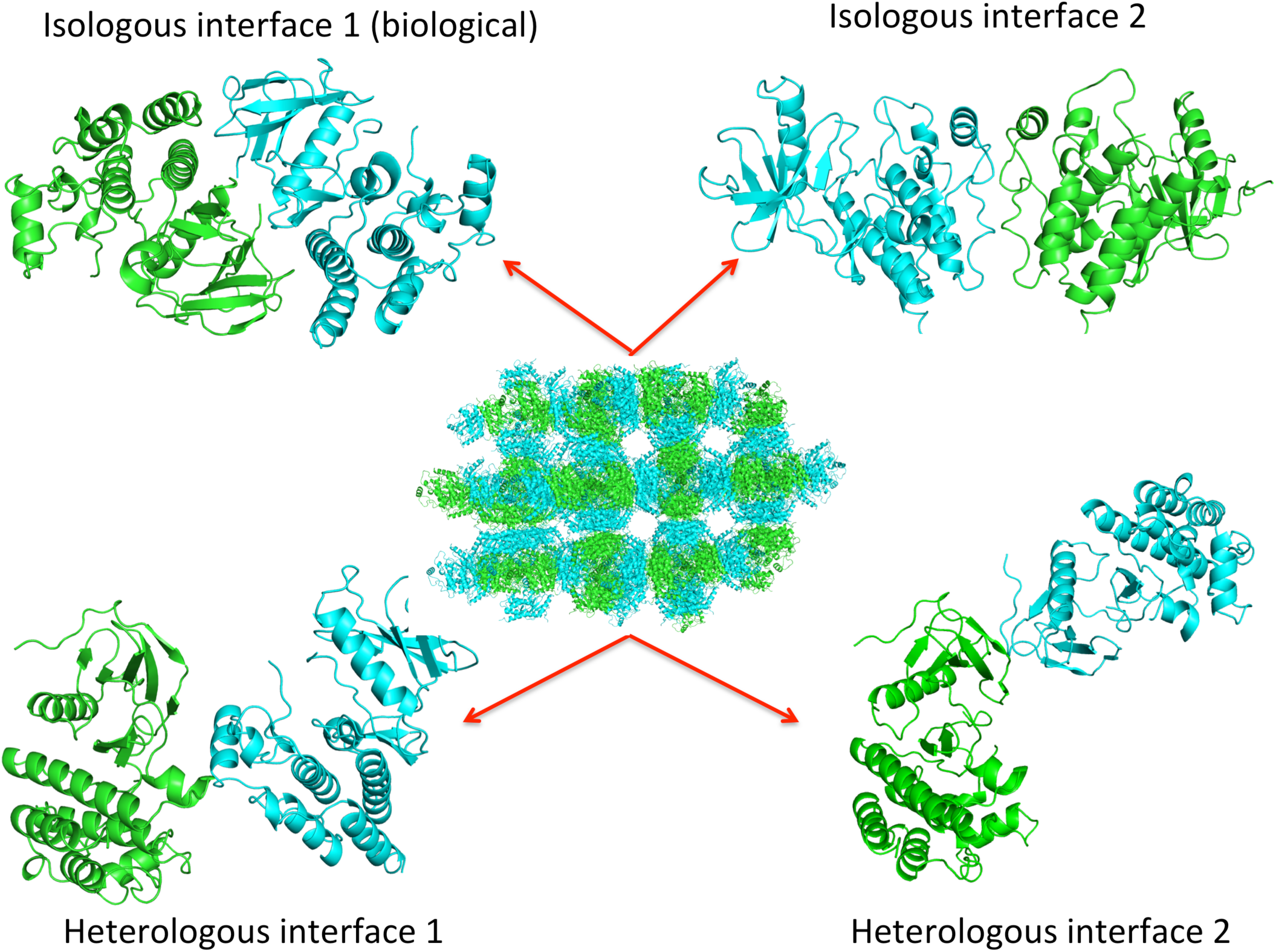
Crystals consist of multiple isologous and heterologous interfaces. The crystal lattice of human BRAF kinase (center) from PDB entry 1UWH contains two isologous interfaces, one of which is the biological dimer (upper left) and two heterologous interfaces.

To study the interfaces in a crystal, it is necessary to consider all interactions within asymmetric units, between asymmetric units within the unit cell, and between asymmetric units in neighboring unit cells. Biologically relevant interfaces may come from any of these sources. Structure files from the PDB contain coordinates for the asymmetric unit and the symmetry operators necessary to build a single unit cell, according to the space group of the crystal. However, some care should be used. Applying these transformations to the asymmetric unit directly sometimes creates asymmetric unit copies in neighboring unit cells, rather than the unit cell containing the original ASU. To solve this problem [5], it is necessary to use the scale matrix in the PDB file to transform the ASU to fractional coordinates. It is then possible to determine which unit cell (±*i*, ±j, ±*k*) the center of mass of the original ASU belongs to. After applying the fractional symmetry operators (e.g., +x,+y,+z+1/2), each copy of the ASU belongs to a particular unit cell where its center of mass resides. It can then be transferred back to the same unit cell of the original ASU by adding integers to the fractional coordinates. Once all symmetry copies of the unit cell have been built, it can be transformed back into Cartesian coordinates in Ångstroms with the inverse of the scale matrix. Once a unit cell has been produced, a set of 3×3×3 unit cells can be constructed by translation of the original unit cell, which is sufficient for identifying all possible interfaces within the crystal.

On average, crystal structures contain 10 different interfaces [15], any of which need to be considered as possibly part of the biological assembly in the crystal. We analyzed the interfaces in 122,340 crystal structures in the PDB for the number of unique interfaces and whether these interfaces were isologous or heterologous (Table 1). The number of unique interfaces types after clustering similar interfaces is somewhat lower (6.72 per crystal) than the total number of interfaces without clustering (9.82). The prevalence of isologous interfaces is about 30%, but this number rises after removing the smallest interfaces; it rises to 44% if only interfaces over 300 Å^2^ are considered. We analyzed likely non-biological interfaces by considering only structures where the ASU was identical to the first biological assembly, and counting only interfaces between ASUs in the crystal (in the same unit cell or neighboring unit cells). A lower fraction of interfaces are now isologous (78%), and the dependence on surface area is lower.

**Table 1.** Isologous and heterologous interfaces in X-ray crystals

|  | ALL |  |  |  | Non-biological interfaces |  |  |  |
| --- | --- | --- | --- | --- | --- | --- | --- | --- |
| Surface Area Cutoff | None | 100 Å<sup>2</sup> | 200 Å<sup>2</sup> | 300 Å<sup>2</sup> | None | 100 Å<sup>2</sup> | 200 Å<sup>2</sup> | 300 Å<sup>2</sup> |
| #Entries | 122340 | 120864 | 120038 | 117487 | 69429 | 68916 | 67209 | 61579 |
| #Unique Interfaces | 823197 | 634633 | 480025 | 352033 | 312696 | 243021 | 179945 | 123568 |
| Interfaces/Crystal | 6.72 | 5.26 | 4.00 | 3.00 | 4.50 | 3.53 | 2.68 | 2.01 |
| Isologous (%) | 30.2 | 33.8 | 38.0 | 44.1 | 21.7 | 23.2 | 25.5 | 29.7 |
| Heterologous (%) | 69.8 | 66.2 | 62.0 | 55.9 | 78.3 | 76.8 | 74.5 | 70.3 |

Non-biological interfaces were identified as the interfaces between asymmetric units in crystal structures with biological assemblies identical to their asymmetric units.

While the biological assemblies deposited by authors are about 80% correct, we can examine the necessity of building out neighboring unit cells of the central unit cell containing the original asymmetric unit by reading the *pdbx_struct_oper_list* records in the mmCIF files for PDB entries, which define the symmetry operators required to build the biological assemblies from the asymmetric unit for each entry. Operators are labeled by their symmetry number (1,2,…,*N*) for *N* operators and which unit cell they belong to, where 1_555 is the original ASU (symmetry operator 1) of the unit cell (555) that contains the asymmetric unit. So for instance, 2_655 is the ASU generated by operator 2 that belongs to the unit cell one cell further along the x-axis of the crystal lattice. About 14% of PDB entries require protein chains from a neighboring unit cell to build at least one of the annotated biological assemblies.

A number of studies have sought to distinguish the properties of interfaces in biological assemblies, sometimes contrasting these properties with those induced solely by crystallization. There are several data sets that have been developed and used repeatedly in this form of analysis. Ponstingl et al. derived a set of 96 monomeric proteins and 76 homodimeric proteins from available annotations in SwissProt and the publications [16]. They used symmetry operators to build a set of 5×5×5 unit cells from which the largest non-biological interface was retained from each crystal for evaluation. Even when considering only the largest non-biological interface, they found smaller surface areas for the non-biological interfaces (distribution mode 400 Å^2^) than the biological interfaces (mode 1200 Å^2^). Including all non-biological interfaces and larger oligomeric assemblies would of course affect the results. The distribution of atom pair types was also distinct for biological versus non-biological interfaces. Using all interfaces, not just the largest non-biological interface, would produce even more distinct distributions of properties of biological and non-biological interfaces. Other more recent studies have found similar results.

Bahadur et al. examined a set of 122 PDB entries annotated as homodimers based on biochemical or biophysical data such as analytical ultracentrifugation [17]. To determine the correct structures for the homodimers, they built all neighboring ASUs of the deposited ASU and identified the largest C2 symmetric interface. They built candidate homodimers for the biological assembly by detecting the largest C2 symmetric interface in each crystal, whether that dimer was in the ASU or between neighboring ASUs or unit cells. They then compared this dimer to the deposited biological dimeric assemblies (where available) and those from the Protein Quaternary Server (PQS) and selected the most likely dimer in the few cases where there were discrepancies. Some ambiguous cases were discarded. To our knowledge, these coordinates are not publicly available and the benchmark assemblies are not all well defined. They found a mean of 1,940 Å for the homodimer surface areas. They studied the hydrophobicity, hydrogen bonding properties, and amino acid composition of the homodimer interfaces and compared them to a set of heterodimeric interfaces collated by the same research group [18].

The sequence conservation of amino acids in homodimeric and heterodimeric interfaces has been studied extensively, frequently being compared to surface residues not known to be involved in such interfaces [19–28,29**]. The results depend on the sequence set used to build the sequence alignment (e.g., the maximum and minimum sequence identity to the target sequence, the process used to remove redundancy), the measures used to quantitate sequence conservation, whether the structures investigated are homodimers, larger homooligomers, or heterodimers and heterooligomers, and which residues are defined as interface residues (sometimes divided into core and rim residues). For instance, Duarte et al. found better distinction between biological from non-biological interfaces by using only closer homologues of the query sequence, careful reduction of sequence redundancy by sequence clustering, and comparison of sequence entropy of non-interface residues with only the core residues of each interface [29**]. These authors carefully curated sets of biological interfaces and non-biological interfaces by checking the primary literature, and choosing biological and non-biological interfaces with similar surface area distributions. They implemented this in the EPPIC program to distinguish biological from non-biological interfaces in crystals.

Besides sequence conservation, conservation of interface structure across multiple structures of the same or homologous proteins has long been used by crystallographers to identify biologically relevant interfaces and assemblies. We clustered all interfaces in crystals of homologous proteins and determined that the number of crystal forms and percentage of available crystal forms that an interface was observed in was strongly correlated with whether that interface was present in biological assemblies or not [5]. We observed this association more clearly when the proteins in the cluster of interfaces contained some member pairs with less than 90% sequence identity, since it has been observed that some proteins commonly form an interface in different crystal forms of identical sequences. The most notorious example of this is a C2 symmetric dimer of T4 lysozyme that is present in 21 crystal forms and 525 PDB entries [30,31**]. We made this form of analysis accessible in a web database called ProtCID (Protein Common Interface Database) [31**]. ProtCID clusters all homologous crystal interfaces of whole protein chains and individual protein domains as identified by Pfam. We build a 3×3×3 set of unit cells to detect unique interfaces in every crystal structure. IBIS (Inferred Biomolecular Interaction server) at NCBI also clusters interfaces of proteins to identify potential biologically relevant interaction sites on proteins [32]. However, IBIS only analyzes interfaces within the deposited asymmetric units and biological assemblies, and does not build a complete unit cell or neighboring unit cells. It thus misses interfaces that ProtCID detects by building a 3×3×3 group of unit cells.

### Symmetries of biological assemblies

There are numerous forms of symmetry in oligomeric protein assemblies [33]. The most common ones are cyclic (Cn) with one n-fold axis of symmetry and dihedral (Dn), with two perpendicular axes of symmetry (one Cn and one C2). Non-symmetric assemblies have C1 symmetry by definition. Other forms of symmetry in biological assemblies include tetrahedral, octahedral, and icosahedral (mostly viruses). Assemblies can also be pseudosymmetric, for instance when homologous proteins with similar folds form cycles or dihedral assemblies. Symmetry can be global, encompassing the entire assembly, or local, covering only a subset of the subunits in the assembly. For instance an A2B heterotrimer may have an isologous (C2) dimer of the A subunits, which then binds a B monomer.

The cyclic forms include symmetric (isologous) dimers (C2), and larger cycles (C3 trimers, C4 tetramers, etc.), built entirely from heterologous interfaces around the cycle (Figure 2A). Additionally some small interfaces may be induced between non-neighboring monomers or complexes across the cycle. The object in the cycle may be a single protein entity, or a complex of two or more proteins. For instance, a heterohexamer of type A3B3 can be a C3 cycle of AB dimers.

**Figure 2.**
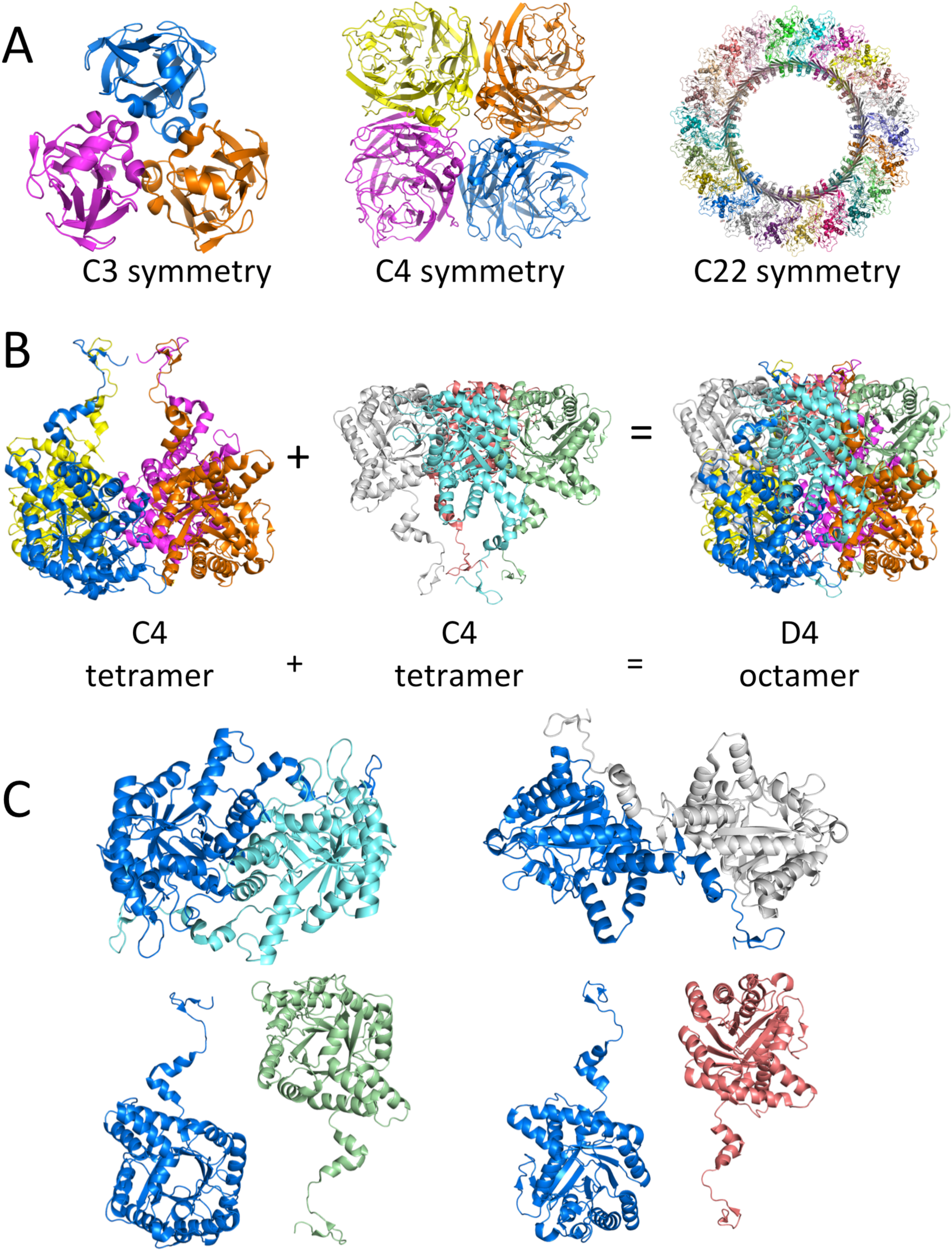
Common symmetries in protein crystals. A. Cyclic assemblies (C3, C4, and C22) consist of heterologous interfaces around the cycle. B. Dihedral assemblies consist of a pair of cyclic assemblies related by a C2 axis (porphobilinogen synthase, PDB entry 3OKB). C. Monomers in one cycle are related by isologous interfaces (or 180° rotation about a common axis even if no contact is made) to those in the other cycle. The relationship between the dark blue monomer in the lower cycle of panel B is shown to the 4 monomers in the upper cycle.

Dihedral (Dn) assemblies can be thought of as a C2 symmetric assembly of two cyclic (Cn) assemblies. For instance, a D4 homooctamer is a symmetric dimer of two C4 homotetramers (Figure 2B). The interfaces between any monomer in one cycle and any monomer in the other cycle are all isologous, as shown in Figure 2C. Alternatively, a D4 homooctamer can be thought of as a cycle of four C2-symmetric homodimers. The interactions between the homodimers are then all heterologous. Heterooligomers can also be viewed with the same symmetry considerations if each subunit is defined as a complex of different molecular species. An A4B4 tetramer can be a C4 cycle of four AB dimers or a D2 assembly of four AB dimers. Some statistics on the distribution of symmetries of biological assemblies from different experimental methods in the current PDB are provided in Table 2. The symmetry of biological assemblies can be determined by the BioJava class QuatSymmetryDetector (https://github.com/biojava/biojava-tutorial/blob/master/structure/symmetry.md) [34], and the program AnAnaS, which not only detects the symmetry but also the symmetry axes [35**].

**Table 2.** Prevalence of assembly symmetry types in the PDB by experimental method

| Symmetry | Diffraction* |  | NMR** |  | Electron microscopy |  |
| --- | --- | --- | --- | --- | --- | --- |
|  | Count | Percent | Count | Percent | Count | Percent |
| C1 | 76,152 | 58.2 | 9,885 | 94.0 | 1,440 | 52.2 |
| C2 | 36,071 | 27.6 | 524 | 5.0 | 245 | 8.9 |
| C3 | 4,534 | 3.5 | 37 | 0.4 | 83 | 3.0 |
| C4 | 1,061 | 0.8 | 20 | 0.2 | 173 | 6.3 |
| C5 | 612 | 0.5 | 10 | 0.2 | 34 | 1.2 |
| C6 | 417 | 0.3 | 1 | 0.01 | 64 | 2.3 |
| Cn>6 | 259 | 0.2 | 1 | 0.01 | 60 | 2.2 |
| D2 | 7,083 | 5.4 | 26 | 0.2 | 18 | 0.7 |
| D3 | 2,156 | 1.6 | 6 | 0.1 | 19 | 0.7 |
| D4 | 674 | 0.5 | 1 | 0.01 | 6 | 0.2 |
| D5 | 235 | 0.2 | - | - | 8 | 0.3 |
| D6 | 123 | 0.1 | - | - | 8 | 0.3 |
| Dn>6 | 135 | 0.1 | - | - | 47 | 1.7 |
| Tetrahedral | 368 | 0.3 | 1 | 0.01 | 11 | 0.4 |
| Octahedral | 348 | 0.3 | - | - | 8 | 0.3 |
| Icosahedral | 394 | 0.3 | - | - | 354 | 12.8 |
| Helical | 188 | 0.1 | 9 | 0.1 | 179 | 12.8 |

\*Diffraction includes X-ray crystallography, electron crystallography, neutron diffraction, fiber diffraction, and powder diffraction
\*\*NMR includes solid-state NMR and solution NMR

While the cyclic and dihedral symmetries have been commonly known since the work of Monod, Changeux, and Wyman [13], the full complexity of symmetric and asymmetric complexes has recently been enumerated by Ahnert et al. in a “periodic table of protein complexes” [36**]. They based this analysis on three steps of oligomer formation consistent with their own experimental data and that of others on protein complex assembly: dimerization, cyclization, and subunit addition. They make an important distinction between “bijective” and “non-bijective” complexes. In the former, every subunit of a particular sequence is in the same topological environment within the complex. In the latter, some subunits are in different environments. This may occur in complexes with uneven stoichiometry amongst subunit types, i.e. those with differing numbers of subunits for different species. It may also occur in asymmetric homodimers, where each subunit uses a different interface to interact with the other. One well known example of this is the asymmetric activating dimer of the kinase domain of EGFR, where the C-terminal domain of one kinase binds to the N-terminal domain of another, activating the latter [37]. This dimer is observed in 14 crystal forms and 104 PDB entries of EGFR, ErbB2, ErbB3, and ErbB4 [38], but only annotated as the biological assembly in 11 entries (Figure 3A).

**Figure 3.**
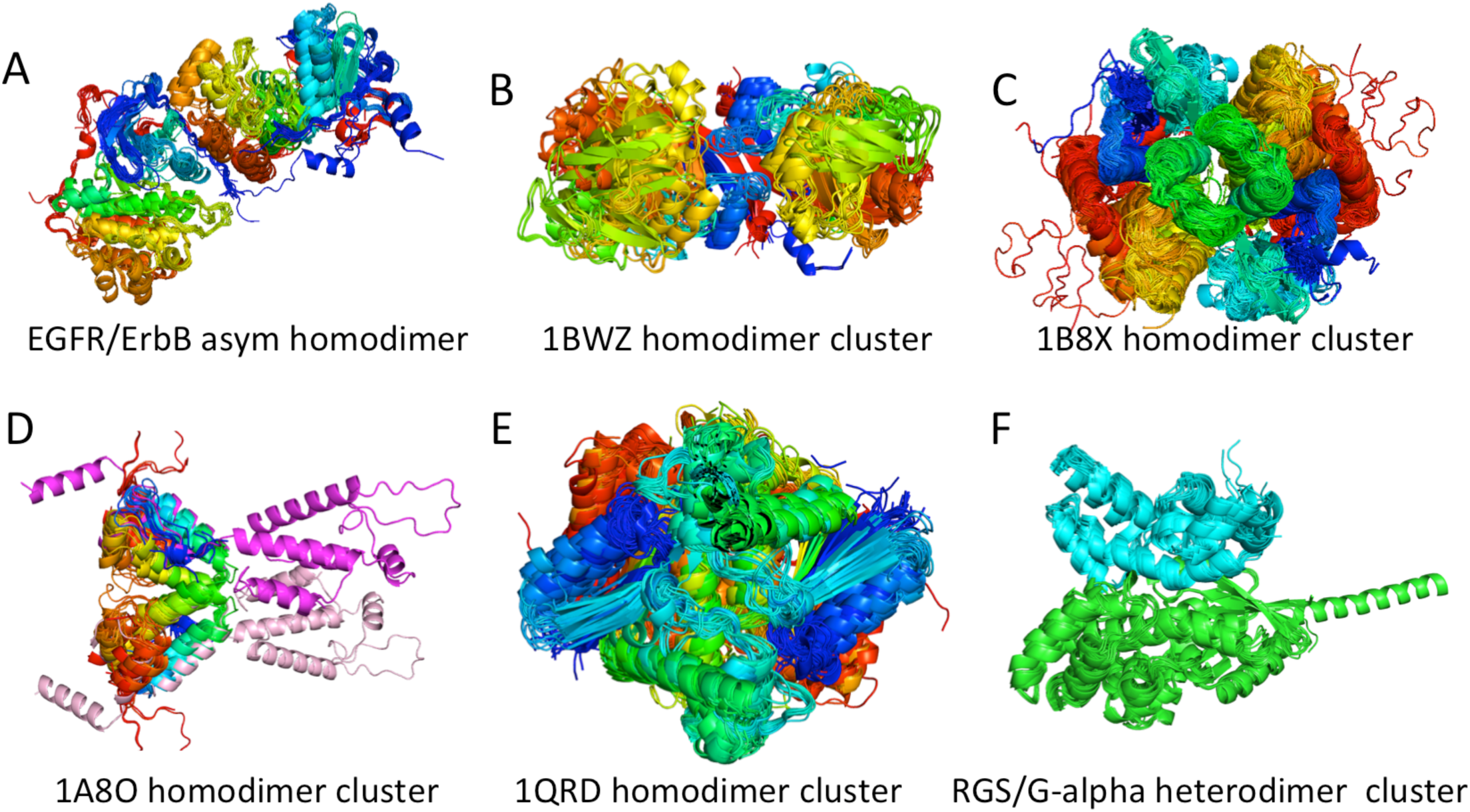
Common interfaces across homologous interfaces in multiple crystal forms. A. The ProtCID cluster of asymmetric homodimers of the EGFR family of kinases (EGFR, ErbB2, ErbB3, ErbB4). B. The ProtCID cluster of DAP epimerases shows that the monomeric annotation for PDB entry 1BWZ in the commonly used Ponstingl benchmark is actually a homodimer. C. The ProtCID cluster of glutathione S-transferases shows the commonly known dimer, including PDB entry 1B8X which is annotated as monomeric in the commonly used Bahadur benchmark. D. HIV capsid proteins form a large assembly with a common dimer in 9 crystal forms of the C-terminal domains including the entry annotated as monomeric in the Ponstingl benchmark (PDB entry 1A8O); the cluster includes an interface from the 18-mer capsid assembly determined by cryo-EM (magenta/pink) (PDB entry 5L93). E. The ProtCID cluster of RGS proteins (regulators of G-protein signaling) and G-protein alpha subunits, all of which have the same interface in the PDB. PISA often fails to identify heterodimeric structures, and predicts a monomer for 9 out of 10 of these structures. F. ProtCID cluster of H-RAS, K-RAS, N-RAS dimers with an interface involving the alpha4 and alpha5 helices from 16 crystal forms.

Some assemblies may be polymeric, such as amyloids, but this may also occur in cytoskeletal proteins, such as actin, and some enzymes. These assemblies can be defined as “helical” even when the relationship between each monomer is simply a translation. The PDB currently contains 408 entries with assemblies annotated as helical; about half of these were determined by cryo-EM and more than half of them have been deposited since 2015. Tsai and Nussinov defined helical symmetries using 4 parameters that are required to define a helical assembly [39**], and their relationship to the asymmetric unit of crystal structures. IMPDH has recently been found to be a filament composed of D4-symmetric octamers [40], which is present in some crystals of IMPDH [41].

It is often assumed that a given protein will have the same form of biological assembly in different crystal structures. Certainly some differences among assemblies in the PDB are due to errors. For example, two crystal structures in different crystal forms may contain the same tetramer but one is annotated as a dimer and the other as the tetramer. In some cases, crystallization conditions or molecular constructs and mutations [42] may interfere with formation of the assembly that is predominant in vivo. The extent to which this occurs in the PDB is unknown, but could be studied with proper assessment of each crystal (building neighboring unit cells, identifying the biological assembly correctly). But many proteins are in equilibrium between assemblies of different sizes, e.g. monomer-dimer or dimer-tetramer equilibria. In most cases, one biological assembly is a sub-complex of the other, e.g. a D2 tetramer and its two constituent C2 dimers. It is not clear how many such cases are represented in the PDB that are truly not related to annotation errors. A few proteins are known to form assemblies with different symmetries, such that one is not a sub-complex of the other. The enzyme porphobilinogen synthase is a D4-symmetric octamer in its active form (Figure 2B), but forms a D3-symmetric hexamer when inactive. This has been demonstrated by Jaffe et al. for the mammalian, plant, and bacterial enzymes [43].

### Benchmarking identification of biologically relevant interfaces and assemblies

Methods have been developed both to distinguish biological from non-biological interfaces within crystals [16,19,28,31,44-46,47**,48-51] and to identify the entire biological assembly within a crystal [6,8,52,53**,54**,55**]. To benchmark these methods properly, the predicted assemblies must be compared to experimentally validated assemblies, not only in stoichiometry and symmetry but also the presence of each interface type. Interfaces and assemblies can be compared either directly by superposition of Cartesian coordinates or indirectly by evaluating the similarity of contacts across each interface. A good benchmark must provide coordinates of the assemblies, and not be limited to stoichiometry and symmetry. It is perfectly possible to identify a C2-symmetric homodimer in a crystal as the biological assembly, when another C2-symmetric homodimer in the crystal is the correct answer.

A number of data sets have been described and used repeatedly as benchmarks for the identification of biological interfaces and assemblies in crystals. Ponstingl et al. compiled a set of 96 monomers and 76 homodimers in the PDB by reference to the published literature and compared the ability of buried surface area and pair interaction scores to predict biological contacts in crystals [16]. Bahadur et al. assembled a set of interfaces consisting of 70 heterodimeric structures, 122 homodimeric structures, and 188 monomeric structures of crystals containing at least one interface with surface area greater than 800 Å^2^ and examined the physical properties of the different interface classes [17,56]. Duarte et al. compiled a set of 78 monomeric structures and a set of 74 oligomeric structures, and consulted the literature to identify the correct assemblies [29**]. After removing duplicate PDB entries, Korkmaz et al. recently combined the Ponstingl, Bahadur, and Duarte sets into a benchmark of 543 PDB entries consisting of 248 monomers, 192 homodimers, 18 heterodimers, 25 homotrimers, 1 ABC heterotrimer, 8 A2B2 heterotetramers, 36 homotetramers, 11 homohexamers, 2 A2B2C2 heterohexamers, 1 A3B3 heterohexamer, and 1 homooctamer [54]. Korkmaz et al. provided the list of PDB entries, the stoichiometry of the biological assembly for each entry (A, A2, AB, A3, A2B2, etc.), and the symmetry of each biological assembly (C1, C2, C3, D2, etc.). They used text mining, comparison of the assemblies of similar sequences, and the PISA and EPPIC3 webservers to evaluate their abilities for predicting stoichiometries and symmetries of biological assemblies for the benchmark.

An analysis of the Korkmaz et al. data set demonstrates some of the challenges, often unacknowledged, of benchmarking the assignment of biological assemblies to PDB entries. We first looked at the sequences and crystal forms of the PDB entries in this benchmark. There are several redundancies where there are two entries in the benchmark that have the same Uniprot sequence and are actually the same crystal form (e.g., 1MOR/1MOQ, 8PRK/1WGJ, 1MKA/1MKB, 1CVU/3PGH, 1EJD/1NAW, 1BC2/2BC2, 232L/256L, 1FEH/3C8Y). We compared the stoichiometries and symmetries of the first biological assemblies given by the PDB to those provided in the benchmark. Only 448 of the 543 PDB biological assemblies (82.5%) agree with the benchmark stoichiometries and symmetries. For 60 of the discrepancies, the benchmark assemblies are labeled monomers while the deposited biological assemblies are asymmetric (C1) homodimers (17 cases), C2 homodimers (37 cases), one heterodimer, and 5 larger assemblies. At least in these cases, what is to be predicted – a monomeric assembly -- is unambiguous. However, 21 C2 homodimers in the benchmark are monomeric in their deposited assemblies and three of them are homotetramers in the PDB. The correct homodimer to compare a predicted biological assembly to is therefore not defined, and may be identified differently by various authors.

It is also likely that some of the benchmark stoichiometries are probably wrong, which can be demonstrated by the much larger number of structures available in the PDB now than when the Ponstingl and Bahadur benchmarks were developed. We used the ProtCID database to identify the largest cluster of interfaces across homologous proteins that each PDB entry in the benchmark belongs to. For protein families that are well represented in the PDB and are homodimers or larger, typically ProtCID will show a large interface cluster containing many different crystal forms and homologous (non-identical) proteins. Usually most but not all of the deposited biological assemblies will include the cluster interface. For instance, PDB entry 1BWZ, is a monomer in the Ponstingl benchmark. However, this entry is a structure of *H. influenza* diaminopimelate epimerase that is found in a ProtCID cluster that includes all 8 crystal forms and 16 PDB entries of the Pfam architecture (DAP_epimerase)_(DAP_epimerase). The cluster contains the *C. glutamicum, B. anthracis, A. baumanni, E. coli,* and *M. tuberculosis* DAP epimerases (Figure 3B). Seven of these 16 structures are annotated as homodimers containing this interface, including the *C. glutamicum, B. anthracis, E. coli,* and *M. tuberculosis* enzymes. Another example is the structure of glutathione S-transferase in PDB entry 1B8X. The protein is listed as a monomer in the Bahadur benchmark, but the ProtCID cluster for this chain includes 1B8X and consists of 80 of 83 crystal forms and 245 PDB entries (Figure 3C), 98% of which are annotated with the ProtCID cluster dimer. GSTs are well known as dimers [57]. PDB entry 1A8O is the HIV capsid C-terminal domain [58], which is a monomer in the Ponstingl benchmark. The same interface in 9 crystal forms (Figure 3D) is also present in a cryo-EM structure of the HIV capsid [59], which demonstrates the potential utility of cryo-EM structures for benchmarking and validating assemblies for X-ray crystal structures. Some other structures in the benchmark that are annotated as monomers but are probably dimers include 1EHY, 1CLU, 1AYI, 1BKZ, and 2YVW. The same is true of 6 benchmark tetramers that are represented by monomeric or dimeric biological assemblies in the PDB.

The situation may actually be worse than this, because it is *not* sufficient to predict the stoichiometry and symmetry of the biological assembly, since crystals can have more than one assembly of the same symmetry. For instance, authors may submit one C2 homodimer as the biological assembly to the PDB but a method may predict that another C2 homodimer in the same crystal is the correct biological assembly. We compared the biological assembly structures with the ASU structures of all entries in the benchmark which had the same symmetry and stoichiometry. We found that for two cases, both the ASU and the first deposited biological assembly were C2 homodimers but they were *different* homodimers (PDB entries 9WGA and 3DA8), and one case where the ASU and the first biological assembly are different A2B2 C2-symmetric heterotetramers (2SCU). It is therefore necessary to compare the actual assembly deposited in the PDB to the predicted assembly. This can be accomplished by structure alignment of the assemblies, taking account of the exchangeability of subunits with the same sequence, or by comparison of interfaces and symmetry of the complexes. This technique was used by Dey et al. to identify biological assemblies by comparison of the deposited biological assemblies across PDB entries [55**].

Recently, Baskaran et al. [47**] developed a very large benchmark of biological and non-biological interfaces to test their previously developed EPPIC program [29**] (described above). They took advantage of our ProtCID database, using stringent criteria to detect biological interface clusters in ProtCID: each cluster they included was present in at least 10 crystal forms and 80% of the crystal forms for the protein family were required to be in the cluster. ProtCID provides all of this information. After removing redundancy and adding 165 dimers from NMR structures, this resulted in a benchmark of 2,831 biological interfaces in their *BioMany* data set. To derive a set of non-biological interfaces, the relied on screw-axis and translation symmetries in crystals, which are typically not part of biological assemblies (except helical filaments which they removed), resulting in 2,913 interfaces in a data set called *XtalMany*. While the Bahadur and Ponstingl benchmarks are still frequently used, we believe that the *BioMany/XtalMany* data set and other sets constructed in a similar fashion are much more valuable for benchmarking biological interface discrimination in crystal structures.

### Methods for identifying biologically relevant interfaces in crystal structures

As described above, many methods have been developed for analyzing which crystal interfaces may be biological and which ones are artifacts of crystallization. We review several of the more recent ones. One important feature of some of the recent work is more rigorous separation of training and testing sets. Many machine learning methods benefit from balanced training data, as we showed for predicting the phenotypes of missense mutations [60]. We also found that balanced accuracy (the average of true positive rate (TPR) and true negative rate (TNR)) was useful as a measure of assessing binary predictions, since it guards against overprediction of either positives or negatives when testing sets are imbalanced. This measure was highest when the training sets for an SVM were balanced, but it was not dependent on whether the testing set was balanced.

Da Silva et al recently developed IChemPic, a random forest method trained on a balanced set of 150 biological interfaces and 150 non-biological interfaces and atom-atom contact features, and tested it on an independent set of 50 biological and 50 non-biological dimers [51]. They calculated all the standard statistics for benchmarking binary predictors (TPR, TNR, PPV, NPV), which is not always done in biological interface prediction. They achieved a balanced accuracy on their testing set of 100 interfaces of 75%, while EPPIC achieved an accuracy of 83% on the same set. Since EPPIC was developed based on some of the same entries in the test set, it cannot be ruled out that this gives EPPIC an unfair advantage on this particular test set. IChemPic outperformed NOXCLASS [45], and PISA [8] by about 3% and outperformed DiMoVo [44] by 21%. The extent of overtraining on the Bahadur and Ponstingl data sets by the earlier methods is evident from the assessment of the same methods on these data sets. Because these benchmarks are not balanced, it is necessary to calculate the balanced accuracy from the sensitivity (TPR) and specificity (TNR) values provided by Da Silva et al. All of the methods (NoxClass, DiMoVo, EPPIC, IChemPic) achieve balanced accuracy rates of 82% to 85% on the Ponstingl and Bahadur data sets (except for a value of 69% for NOXClass on the Bahadur data set). The IChemPic method achieved similar values for TPR and TNR, while the other methods often achieved quite different values, which is an advantage for IChemPIC.

Liu et al. used B-factors of interface residues, and showed that biological interfaces have lower B-factors on average than crystal contacts [49]. They did not provide TPR values, but rather Matthews correlation coefficients for comparing their results to EPPIC and PISA on the Bahadur and Ponstingl data sets. Their MCC rates were slightly higher but it is difficult to tell if their tested method was trained in part on the Bahadur and Ponstingl data sets. Luo et al. recently used packing in interfaces to train a random forest model, and claimed slightly higher balanced accuracies than PISA and EPPIC, although each method was tested on different data sets (those which they were not trained on) [50].

Very recently, Yueh et al. used a novel approach of using docking software to determine whether the ability to dock two monomers together to form a biological dimer and the failure to do so for non-biological dimers might be used to identify biological from non-biological interfaces in crystals [61**]. This method, ClusProDC which uses the ClusPro program for docking [62], performed about as well as EPPIC and somewhat better than PISA.

### Methods for identifying biologically relevant assemblies in crystal structures

There are fewer methods for identifying the likely biological assembly [6,8,52,53**,54**,55**] in a protein crystal than there methods for identifying biologically relevant interfaces. There are several algorithmic features that these methods should implement, although not all do. First, they should build the asymmetric units of the entire unit cell and then neighboring unit cells. If the unit cells are built properly (i.e. each ASU copy in the first unit cell is built by applying a symmetry operator and then translated back to the unit cell of the deposited ASU), then only a set of 3×3×3 unit cells needs to be built. Second, interfaces are evaluated by surface area, biophysical properties, symmetry (i.e., if homodimeric, isologous or heterologous), sequence conservation, and any other properties that might indicate participation in an assembly; they should also be clustered so that similar interfaces are grouped together (e.g., each copy of the same homodimer interface that might appear in a tetrameric ASU). Fourth, explicitly or implicitly, it is usually assumed that the entire crystal is made of biological assemblies with the same stoichiometry, interfaces, and symmetry; that this assembly covers the whole crystal; that every interface of a given type is engaged, if present in a biological assembly, or not engaged, if considered a crystal contact; and that each assembly is closed, i.e., it does not extend the full length of the crystal in any dimension. These rules apply to most crystals but there are known exceptions which have to be considered. Finally, the possible assemblies are evaluated for their biological relevance using the properties of the interfaces or the whole assembly already described, and one or more sets of Cartesian coordinates for the assemblies are produced.

One of the earliest programs (1998) for producing biological assembly coordinates from crystals was the Protein Quaternary Server (PQS) of Henrick and Thornton [52]. PQS built all ASU copies in a 3×3×3 group of unit cells, and starting with a monomer in the asymmetric unit, proceeded to add monomers that have interfaces considered likely to be biological (by surface area, number of contacts vs length of chain, etc.) The final possible assemblies were then evaluated by change in surface area from the full assembly to dissociation into monomers with a cutoff of at least 400 Å^2^ for each monomer in a valid assembly as well as desolvation properties, hydrogen bonds, salt bridges, and disulfide bonds among the monomers in the assembly. One interesting example they gave is PDB entry 1QRD, which is a C2-symmetric homodimer in the ASU, but PQS generated a different C2-symmetric dimer as the correct assembly, even when the annotation at the time was monomeric. ProtCID shows that all 38 crystal forms currently available of quinone reductases contain the PQS-generated dimer, and 97% of them are annotated as the biological assembly.

In 2003, Ponstingl, Kabir, and Thornton improved on PQS, first by developing a benchmark of monomers, dimers, trimers, tetramers, and hexamers, restricting the generated oligomers to point group symmetry (so that every monomer in the assembly formed the same type(s) and number(s) of interface) [6]. They built up assemblies, as PQS does, forbidding translation symmetry and screw-axis operators in the assembly, and then partitioning them until valid (Cn or Dn) assemblies with high stability scores were formed. The accuracy on their benchmark was 84%, but the score cutoff value for partitioning assemblies was set on the same data set (i.e., possible overtraining). The method was implemented in a program called PITA, although this method is no longer available online.

Krissinel and Henrick developed the PISA method in 2007, emphasizing the necessity of considering the thermodynamics of the formation of entire assemblies (rather than the sum over interfaces) and an evaluation of the chemical thermodynamics of assembly formation including entropy and desolvation [8]. They did not assume that a crystal comprises only identical assemblies, explicitly allowing for an assembly to be crystallized with some of its dissociated components. Each interface type is either engaged or disengaged. The algorithm generates all possible closed assemblies using a backtracking algorithm to evaluate all 2^*N*^ assemblies that can be generated for *N* interface types (i.e. each interface type is either engaged or not engated, and so there are 2^*N*^ possible assemblies. Branches that involve parallel monomers may be cut, and interfaces that are induced by other interfaces can be pruned from the search space (e.g., if formation of AB and BC interfaces induces AC, then AC can be removed from the list of possible interfaces). Ultimately, PISA uses three criteria to rank the generated assemblies: 1) larger assemblies are preferred over smaller ones; 2) solutions in which individual assemblies cover the crystal are preferred over solutions with multiple assemblies with different stoichiometries, interfaces, or symmetries; 3) assemblies with higher free energies of association are preferred over those with lower values. PISA was perhaps the first program for biological assemblies to consider the complexities of ligands and DNA binding. The parameters of PISA were trained on the Ponstingl data set [6] such that PISA predicted an assembly of the right size (disregarding whether the predicted assembly contained the right interfaces or symmetry). PISA could be profitably retrained now on a much larger data set that can now be constructed. In our experience, PISA often fails to score a heteromeric assembly as the top assembly in crystals with more than one type of subunit. This often occurs when one of the subunits is small, such as a peptide or small protein domain. For example, complexes of proteins with the Pfams (G-alpha) and (RGS) are found in 10 crystal forms; all of these crystal forms contain a common heterodimer in ProtCID that is annotated by the authors as the biological assembly in all 10 PDB entries (Figure 3E), but PISA produces monomeric predictions for 9 out of these 10 entries.

In 2011, Mitra and Pal utilized a set of 10 features of interfaces – based on packing, surface complementarity, surface area, and solvation energies – with a naïve Bayes classifier model to predict full quaternary structures in crystal structures [63]. They build symmetry copies of the ASU, and build up “functional units” with their classifier for biological and non-biological interfaces. Assemblies larger than monomers that do not have point symmetry are discarded. They focused particularly on the prediction of weak, transient oligomeric structures, which are poorly predicted by most methods. Their training data came from the PiQSi database, which is a manually curated data set of about 15,000 biological assemblies by reference to the published literature and homologous structures [9]. In particular, their method called IPAC performed significantly better on heterodimers than PISA, which as noted performs poorly on heteromeric structures.

EPPIC3, developed in 2018 by Bliven et al., rigorously implements rules for the possible assemblies that can be found in a crystal – building neighboring unit cells, closed assemblies with point group symmetry, and full coverage of the crystal with a single type of assembly [53**]. Each assembly is then evaluated by its conservation scores produced by EPPIC [29**]. Enumerating all of the possible 2^*N*^ assemblies is time consuming. As noted earlier, the tree can be pruned if an interface violates one of the assembly rules. EPPIC3 merges some heteromeric interfaces as a way to speed up the algorithm. One ambiguous feature in EPPIC3 is the handling of heteromeric assemblies; the paper states that interfaces between different sequence identities are merged “in a greedy manner,” and the resulting dimers are used as single entities. A very useful testing set of 1,481 biological assemblies was compiled by clustering sequences in the PDB at 70% sequence identity and choosing one representative from each cluster that had at least three structures and identical assembly annotations (it is not clear if this means only stoichiometry and symmetry (provided by the PDB) or that it also includes comparison of interfaces and coordinates). They showed that EPPIC3 performs slightly better than PISA (85.3% vs 83.5% on their biological assembly data set) with a tradeoff of predicting fewer large assemblies correctly than PISA, which fails more frequently on smaller assemblies than EPPIC3. However, it should be noted that predictions were compared at the level of stoichiometry (assembly size) and not by interfaces or symmetry. As noted earlier, it is perfectly possible to predict the right stoichiometry and even the right symmetry but have the wrong biological assembly with incorrect interfaces.

The power of comparing biological assemblies across crystal forms, similar to our comparison of interfaces in ProtCID, has been shown recently by several groups [54**,55**,64**,65]. Dey et al. compared biological assemblies annotated in the PDB and those from PISA and EPPIC2 interfaces across groups of homologous proteins in order to determine whether conservation of biological assemblies might be used to annotate previously defined biological assemblies for each PDB entry as correct or incorrect [55**]. They properly compared not only stoichiometries and symmetries but also performed superpositions of assemblies of the same size to determine whether two assemblies were the same. This has to be done carefully, since the order of monomers in a homooligomeric assembly can be in different order in one structure than another. For symmetric assemblies, every monomer is in the same topological environment as any other, so superposition of just one monomer between the assemblies will serve to identify the correct order. The identification of similar assemblies across entries was referred to as QSalign. Monomers could be identified by the lack of similar assemblies across PDB entries in a family, which they implemented in a strategy called anti-QSalign. Together QSalign and anti-QSalign form a procedure, QSbio, for determining biological assemblies from comparison of homologous proteins. Assemblies are deemed “correct” if they are found in 2 or more structures that have less than 80% sequence identity, with larger assemblies considered before smaller ones. An assembly that is 100% identical in sequence to a “correct” assembly (but does not contain the same assembly) is labeled as “ambiguous” if it is larger than the “correct” assembly, and incorrect if it is the same size or smaller. QSbio is limited by the biological assemblies in the PDB and generated by PISA, so while it may detect that an assembly is incorrect because all other structures all have the same assembly as each other, it does not generate the assembly for the incorrect entry, even when the “correct” assembly may be present in the crystal of the incorrect entry. For weak or transient interactions, it is possible that the correct assembly has not been identified (or at least not for more than a pair of homologous sequences with <80% sequence identity) by any of the authors or PISA, and QSbio would not identify these.

Korkmaz et al. recently used a similar procedure to correct the biological assembly annotations in the PDB [54] by comparing assemblies across homologous PDB entries at the level of stoichiometry and symmetry and the annotations from author annotations, PISA, EPPIC3, and a text mining approach of the literature. However, unlike QSbio, they did not perform structural comparisons of the interfaces or coordinates of each assembly, which limits this approach. Bertoni et al. have used a similar approach in SwissModel by comparing biological assemblies of different PDB entries in a family of potential templates for building homology models of oligomeric complexes [64**]. They compared the interfaces of oligomers to determine whether assemblies were isomorphous using a variant of the Q-score we developed for interface comparison in ProtCID. Similarly, Sudha and Srinivasan compared the stoichiometries, interfaces, and symmetries of distantly related assemblies of proteins in different PDB entries, and correlated similarities and differences to functional states [65].

### There are exceptions to every rule

There are several forms of biological assemblies that are not necessarily well handled by the methods described above, which have implemented rules that define the valid assemblies from crystals. These rules probably prevent more incorrect predictions than they forbid correct predictions that violate the rules. But nevertheless, the violations are often very interesting and in some cases unexpected or undetected by authors. These situations include complexes that have uneven stoichiometry, asymmetric configurations (even in homodimers and larger oligomers), and filamentous structures with helical symmetry. But there are other rules that also may be broken. For instance, it is not clear whether every crystal is constructed of only one assembly type. As Krissinel has pointed out, crystal formation of weaker dimers may occur such that both the monomer and dimeric molecules crystallize together. The extent of this phenomenon is not known. Some of the earliest structures of p53 with DNA contained three copies of p53 in the asymmetric unit (PDB entries 1TUP and 1TSR), only two of which interacted with the DNA in a sequence-specific fashion [66]. This crystal form can be considered a superassembly of the dimer/DNA complex and a third p53 monomer.

The challenge of identifying weak complexes, especially transient homodimers, remains [67]. One way to do this may be to examine multiple crystals of the same protein or homologous proteins. In this way, we have identified a C2-symmetric homodimer of H-RAS, K-RAS, and N-RAS in a ProtCID cluster that contains 16 crystal forms (Figure 3F). There is experimental evidence published recently that this alpha4-alpha5 dimer is at least one of the biologically relevant dimers of RAS proteins [11,68,69]. EPPIC3, PISA, and QSbio all have trouble identifying this dimer.

### Conclusions

The several methods that depend on comparing assemblies and interfaces in different crystals of homologous proteins will only improve as the number of structures in the PDB increases. Methods that depend on sequence conservation will also improve as the sequence databases increase in size, especially with the addition of more eukaryotic genomes. The rapid rise of cryo-electron microscopy and the vastly improved resolution of these structures provide new opportunities in the annotation of biological assemblies crystal structures of homologous proteins. More than half of cryo-EM structures have at least two copies of at least one subunit type, and therefore provide non-crystallographic information on homooligomeric interfaces. Finally, the accurate determination of biological assemblies also has a significant impact on using comparative modeling of assemblies [64**,70-75], since these methods depend on the accurate annotation of PDB entries used for template-based modeling.

